# Sero-epidemiological investigation and risk factors associated with camelpox in pastoral areas of Somali region, eastern Ethiopia: a cross-sectional study

**DOI:** 10.1101/2024.06.05.597649

**Authors:** Hassan Abdi Arog, Abdullahi Adan Ahad, Haben Fesseha

## Abstract

Camelpox is a highly significant viral disease that has a major economic impact on camels in Ethiopia. However, the epidemiology of the disease in the country, particularly in Somali region, is currently not well understood. The objective of this study was to estimate the prevalence of camelpox and identify associated risk factors in order to implement effective disease control measures in the study area. A cross-sectional design was employed from January 2023 to July 2023, involving a sample of 374 camels from 75 households in two districts and six peasant associations (PAs). Serum samples were tested using a competitive enzyme immunoassay (c-ELISA) to detect camelpox-specific antibodies. The overall seroprevalence of camelpox infection in the study area was found to be 16.0%. We administered structured questionnaires to camel owners to gather additional information on potential risk factors. Variables such as age, sex, and seasonal patterns were found to have a significant association with camelpox seropositivity. Female camels exhibited 3.2 times higher odds of infection compared to male camels, while young dromedaries aged between 6 months and 4 years were found to have a 2.3 times higher risk of infection than adults, indicating susceptibility to the age factor. Furthermore, the risk of infection was found to be 26 times higher during the rainy season than to the dry period. Thus, by identifying contributing factors, effective preventative measures, such as an appropriate vaccination strategy, can be developed to reduce the spread of camelpox and the associated economic losses. This study provides valuable insights for disease control and management practices.

## Introduction

The global camel population is estimated to be more than 35 million heads, where one-humped camels (*Camelus dromedarius*) occupies the largest camel distribution [1, 2]. Camels are vital for meat and milk production in many regions, particularly in Africa and Asia. The one-humped camel makes up around 95% of the total population of Old-World camels [3]. Africa is home to over 80% of the world’s camel population, with 60% of these camels residing in the Horn of Africa, including Somalia, Sudan, Ethiopia, Kenya, Djibouti, and Eritrea. The proportion of one-humped camels is higher in this region compared to the rest of the world [4]. In Ethiopia, the camel population is estimated to be over 8.1 million, placing the country third in Africa [5]. Camels are mainly found in the arid and semi-arid lowlands of the Somali, Afar, and Southern Oromia regional states, where nomadic herders make up the majority of the population [5, 6]. Camels are highly valued in the region and play a crucial role in milk and meat production, transportation, and trade [4]. These camels are bred to withstand the harsh conditions of the arid and semi-arid rangelands, where pastoralism is the primary way of life. However, one of the major challenges faced by the pastoral community in the region is infectious camel diseases, including camelpox [7].

According to the World Organization for Animal Health (OIE), camelpox is an acute and highly contagious viral disease that primarily affects camels. It belongs to the Orthopoxvirus genus of the Poxviridae family [8]. Camelpox causes significant economic losses and poses a potential public health threat. It is characterized by the formation of nodular skin lesions, fever, and general signs of septicemia. The disease can be transmitted to other susceptible species through close contact with infected animals [9].

In Ethiopia, camelpox has been increasingly reported in various regions of the country [10]. For example, in the Borana Zone of the Oromia region, the prevalence of camelpox was 0% during the dry season and 14.2% during the minor rainy season based on clinical diagnosis [11]. In the Amibara and Awash Fentale zones of Afar, the overall seroprevalence was found to be 21.6% and 16.7%, respectively [12]. Additionally, the first molecular detection of camelpox virus in the Afar region was reported by Gelagay *et al*. [13]. However, to the best of our knowledge, no previous research has investigated the epidemiology and risk factors associated with camelpox occurrence in the lowland area of the Somali region in eastern Ethiopia, where camelpox circulation is possible due to geographical differences. Therefore, the main objective of this study was to estimate the prevalence status and identify potential risk factors contributing to the spread of camelpox among susceptible hosts in the study area. The findings will help policy makers to implement effective control measures to reduce the disease burden among ground pastoralists.

## Materials and methods

### Description of the study area

This study was conducted in two randomly selected districts of the Jarar zone, Somali Region: Degahbour and Gunagado (Fig 1). The Jarar zone consists of a total of 11 districts, with Degahbour and Gunagado falling under its administrative jurisdiction. Degahbour district is located 160 kilometers away from Jigjiga city, the capital city of the Somali Region. It is also situated 630 kilometers away from Addis Ababa and 103 kilometers away from Harar. The district has an elevation of 1,044 meters above sea level at a latitude of 8° 13’ N and a longitude of 43° 34’ E. which is known for its significant camel population, and experiences high temperatures and limited rainfall, resulting in a dry and semi-arid environment. Camels play a vital role in the livelihoods and culture of the people in the area, with an estimated camel population of around 116,790 in this particular district [14].

**Fig 1.**
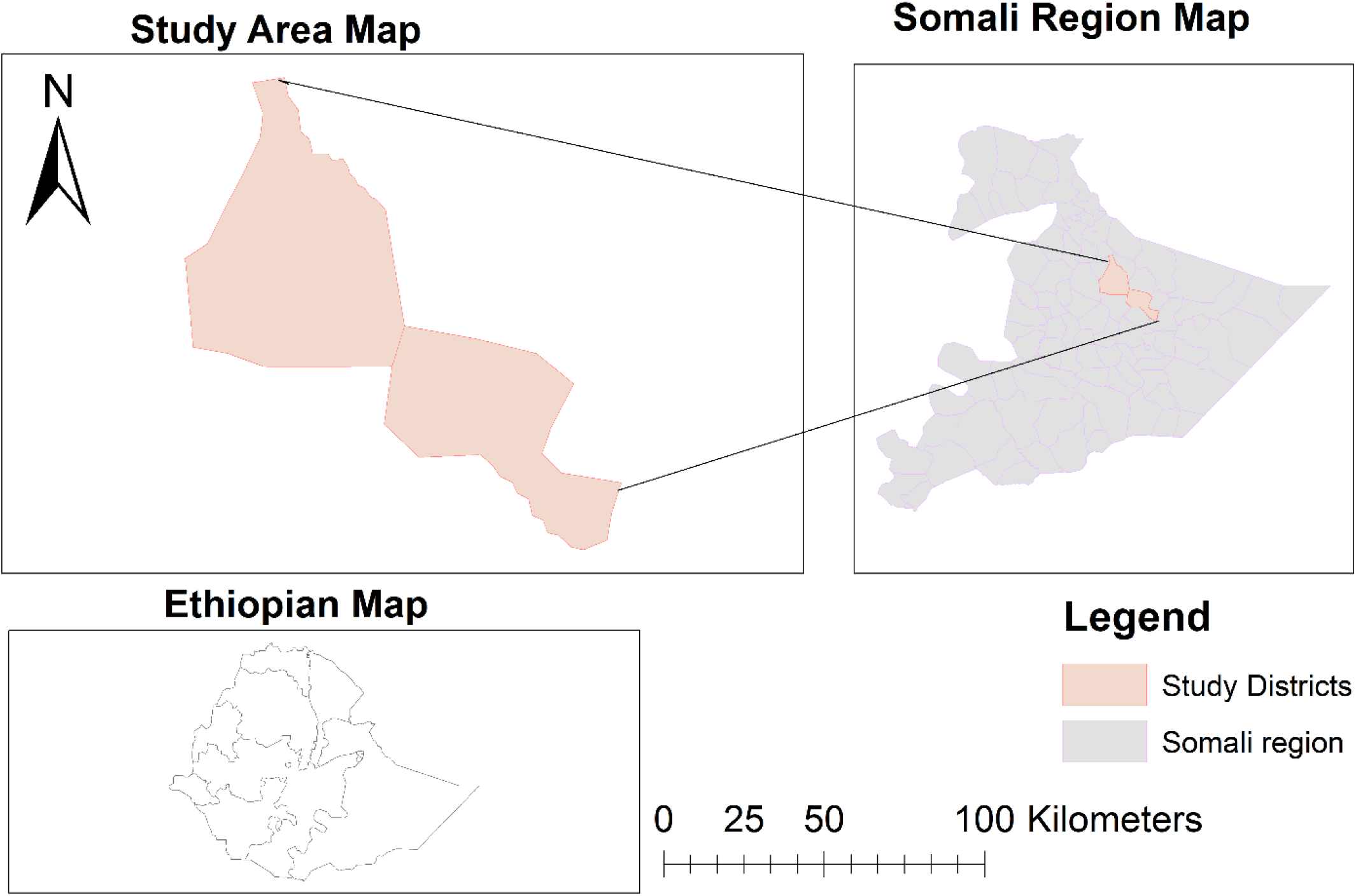
Map of the study areas extracted from an Ethiopian shapefile using QGIS version 3.20.0.

Geographically, Gunagado district is located in the southeastern part of the city and covers a large area, with an elevation of 1,640 meters above sea level. It experiences two main rainy seasons: *Gu’* (spring) and *Deyr* (fall). The spring rainy season lasts from March to May, while the fall rainy season extends from September to November. The average annual temperature in Gunagado district is 29°C, with an annual rainfall ranging from 400 to 600 mm. The predominant mode of production in this area is an extensive system, where over 90% of the population moves from one place to another in search of pasture and water, and the approximate number of camels is 137,521 [15].

### Study population

The study animals in this research were domestic dromedary camels that were raised by Somali camel owners in selected districts of Jarar Zone. The country has a total of 8.1 million camels (CSA, 2020), and the estimated number of camels in the Somali region is approximately 4.5 million where Jarar Zone alone has around 258,477 camels, making it the most densely populated region in terms of camels which were considered as the target population for this study [16]. The ages of the camels were classified as young (6 months to 4 years) and adults (older than 4 years), and the sex of the camels was categorized as female and male. Additionally, the seasons were divided into dry and wet seasons [11, 17].

### Inclusion and exclusion criteria

To minimize the influence of herd migration bias, only animals that were part of the study were selected for sampling from specific districts in the Jarar Zone, Somali Region. The study included male and female camels that were at least 6 months old. An exclusion criterion was used to remove camels from the sampling group if they had been vaccinated against camelpox in the past year, as confirmed by historical information from the camel owners. This exclusion was implemented to avoid any potential interference from vaccination-induced antibody development, which could impact the natural infection.

### Study design and sampling methods

From January 2023 to July 2023, we conducted a cross-sectional study to investigate the seroprevalence of camelpox and identify potential risk factors associated with the occurrence of the disease in select districts in the Jara zone of the Somali Region of Ethiopia. We employed a purposive sampling technique for this study. We chose the Jarar zone as our study area and selected two districts, Degahbour and Gunagado, based on factors such as camel population density, geographical location, accessibility and convenience.

To collect samples, we used a simple random sampling method. This allowed us to select peasant associations (PAs), 75 households, and study camels in a way that ensured representative sampling. In total, we selected six PAs for the study: three from each district. In Degahbour, we chose Bulale, Dogorjir, and Dabile as the PAs, while in Gunagado, our selected PAs were Balisaredo, Bulhan, and Kararo. To gather data on potential risk factors, we utilized a structured questionnaire. The variables recorded included sex, age, season, study sites, and herd size.

### Sample size determination

The sample size estimation for an epidemiological study using a serological assay was performed based on the formula provided by Thrusfield [18]. This calculation took into account a confidence level of 95%, an expected prevalence of 14.2% as reported by Megerssa [11], and an absolute precision of 5%.

Where,

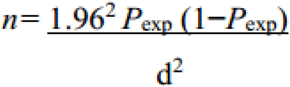

n =required sample size

Pexp=expected prevalence

d_2_=desired absolute precision (5%)

The total sample size was determined by performing calculations, resulting in a sample size of 187. However, to enhance the representativeness and accuracy of the sample, a design effect of 2 was considered. Therefore, a minimum sample size of 374 serum samples was required.

## Data collection

### Serum sample collection

Based on a study of animals, a total of 10 ml of blood was collected from camels’ jugular veins using sterile, plain vacutainer tubes. The blood tubes were then left undisturbed at room temperature overnight. As a final step, the collected serum samples were transferred into sterile cryogenic tubes. The specimens were transported in an icebox to the Jigjiga Regional Laboratory and Diagnostic Center (JRLDC) and stored at -20 °C in a deep freezer until they were ready for laboratory analysis. The analysis aimed to detect camelpox antibodies using the sandwich enzyme-linked immunosorbent assay (ELISA) method [19].

### Laboratory analysis

Serum samples collected from camels were analyzed using a sandwich ELISA test, which is a recommended method for detecting camelpox virus-specific antibodies, according to Khalafalla *et al*. [20]. This method is endorsed by the Office International des Epizootics.

This sandwich ELISA test is a widely used immunoassay technique. In this assay, an antibody specific to camelpox is pre-coated onto a 96-well plate. Controls, test samples, and horseradish peroxidase (HRP)-conjugated reagents are added to the wells and incubated. After incubation, unbound conjugates are removed by washing the plate with a wash buffer. To quantify the HRP enzymatic reaction, a TMB substrate is added to each well. Only wells that contain sufficient camelpox will produce a blue-colored product, which later changes to yellow upon addition of the acidic stop solution. The intensity of the yellow color is directly proportional to the amount of camelpox bound on the plate. The optical density (OD) of the yellow color is measured spectrophotometrically at 450 nm using a microplate reader. By comparing the OD values of the test samples to the controls, the presence of camelpox can be determined. The entire test procedure was conducted at the Jigjiga Regional Laboratory Diagnostic Center (JRLDC). The details of the test principle, materials and reagents used, the test procedure, and interpretation of the results were described by Aboul *et al*., [21, 22].

### Ethical approval

The study was conducted in accordance with the guidelines of the Declaration of Helsinki. Approval for this study was granted by the Jigjiga University Review Board, college of veterinary medicine. The committee reviewed the study in accordance with the university policy environment safeguarding animal rights and welfare and the committee granted approval with permit (No. JJU/REC/002/2023). An oral consent was also obtained from camel herders.

### Data management and analysis

Data collected in the field was recorded and stored in Epi data entry version 3.0. The data was then imported into STATA® version 14.0 statistical software to calculate disease prevalence and odds ratios. Logistic regression analysis was used to measure the association between potential risk factors and the occurrence of camelpox seroprevalence. First, univariable analysis was performed on all predictors by fitting them into separate logistic regression models to assess their unconditional associations. All variables with a p-value of less than 0.25 were included in the multivariable analysis to reduce the number of predictor variables with unconditional association. Finally, odds ratios and the 95% confidence interval (CI) were calculated, and disease-associated risk factors with a p-value of less than 0.05 were considered significant [23].

## Results

### Seroprevalence of camelpox seropositivity

The results of this study showed that out of the 374 dromedary camels tested using a sandwich ELISA test, 60 tested positive for camelpox infection. Therefore, it was determined that the overall seroprevalence of camelpox disease in the two selected districts of Jarar Zone in Somali Regional State was 16.0% (95% CI: 12.0–20.0). This study found that Gunagado district had a relatively high prevalence of 19% (95% CI: 14.0–25.0), while Degahbour district had a prevalence of 12% (95% CI: 7.0–17.0). However, the difference between the two districts was not statistically significant.

### Risk factors associated with camelpox

The study revealed that young adults had a significantly higher prevalence of camelpox antibodies, at 21% (95% CI: 15.0–26.0), compared to adults in other age groups, who had a prevalence of 11% (95% CI: 6.0–16.0). Additionally, when considering gender, female dromedaries showed a higher serological prevalence, at 21% (95% CI: 16.0–27.0), than males, at 8% (95% CI: 4.0–13.0), and this difference was found to be statistically significant (p<0.05).

Meanwhile, there was no statistically significant difference in camelpox seropositivity among the three herd sizes (p > 0.05). However, large herds (>50 camels) had the highest prevalence rate at 20% (95% CI: 14.0–26.0), followed by small herds (<25) at 16% (95% CI: 9.0–23.0) and medium herds (>25 and 50 camels) at 10% (95% CI: 4.0–15.0).

Regarding the district analysis, Gunagado district had a higher infection rate than Deegahbour district. Nevertheless, the difference was not statistically significant (P > 0.05). Similarly, this study showed that the disease occurrence was higher during the wet season than the dry (winter) period, and it was statistically significant (p < 0.05). The table below (Table 1) shows hypothesized risk factors considered, such as age groups, sex, season, herd size, and study districts associated with seropositive camels.

**Table 1.** This table presents the results of a univariate analysis that evaluates the risk factors associated with camelpox seropositivity.

| Variables | Categories | Sera tested | Sera positive | Prevalence %<br>(95% CI) | P-value |
| --- | --- | --- | --- | --- | --- |
| Districts | Degahbour | 166 | 20 | 12 (7.0-17.0) | 0.06 |
|  | Gunagado | 208 | 40 | 19 (14-25) |  |
| Sex | Male | 155 | 13 | 8 (4.0-13.0) | 0.001* |
|  | Female | 219 | 47 | 21 (16.0-27.0) |  |
| Age | Young | 193 | 40 | 21 (15.0-26.0) | 0.01* |
|  | Adult | 181 | 20 | 11 (6.0-16.0) |  |
| Season | Dry | 158 | 2 | 1 (0.0-3.0) | 0.0001* |
|  | Wet | 216 | 58 | 27 (21.0-33.0) |  |
| Herd size | Small | 100 | 16 | 16 (9.0-23.0) | 0.08 |
|  | Medium | 104 | 10 | 10 (4.0-15.0) |  |
|  | Large | 170 | 34 | 20 (14.0-26.0) |  |

\*Statistically significant

To investigate the main risk factors associated with camelpox seropositivity, a univariable analysis was conducted. This analysis identified several potential factors, including season, age group, sex, herd size, and study districts, based on their significance threshold (P≤0.25). Subsequently, the risk factors underwent a multivariable logistic regression analysis to further investigate their impact.

The results of the multivariable logistic regression analysis showed that age, season, and sex were the only variables significantly associated with the occurrence of seropositivity to camelpox (p < 0.05). Young dromedaries had a higher risk of camelpox infection compared to adult dromedaries, with an odds ratio (OR) of 2.3 (95% CI: 1.2–4.2). The statistical analysis revealed a significant association between age and the occurrence of camelpox (p < 0.05).

The odds ratio (OR) for camelpox seropositivity in females was 3.2 (95% CI: 1.6-6.4), indicating that females were 3.2 times more likely to contract the disease compared to males. The association between sex and camelpox occurrence was found to be statistically significant (p < 0.05). Furthermore, the study revealed that camelpox was significantly more prevalent during the rainy season compared to the dry season. According to Table 2, the likelihood of camelpox occurring during the rainy season was 26 times higher than during the dry season.

**Table 2.**
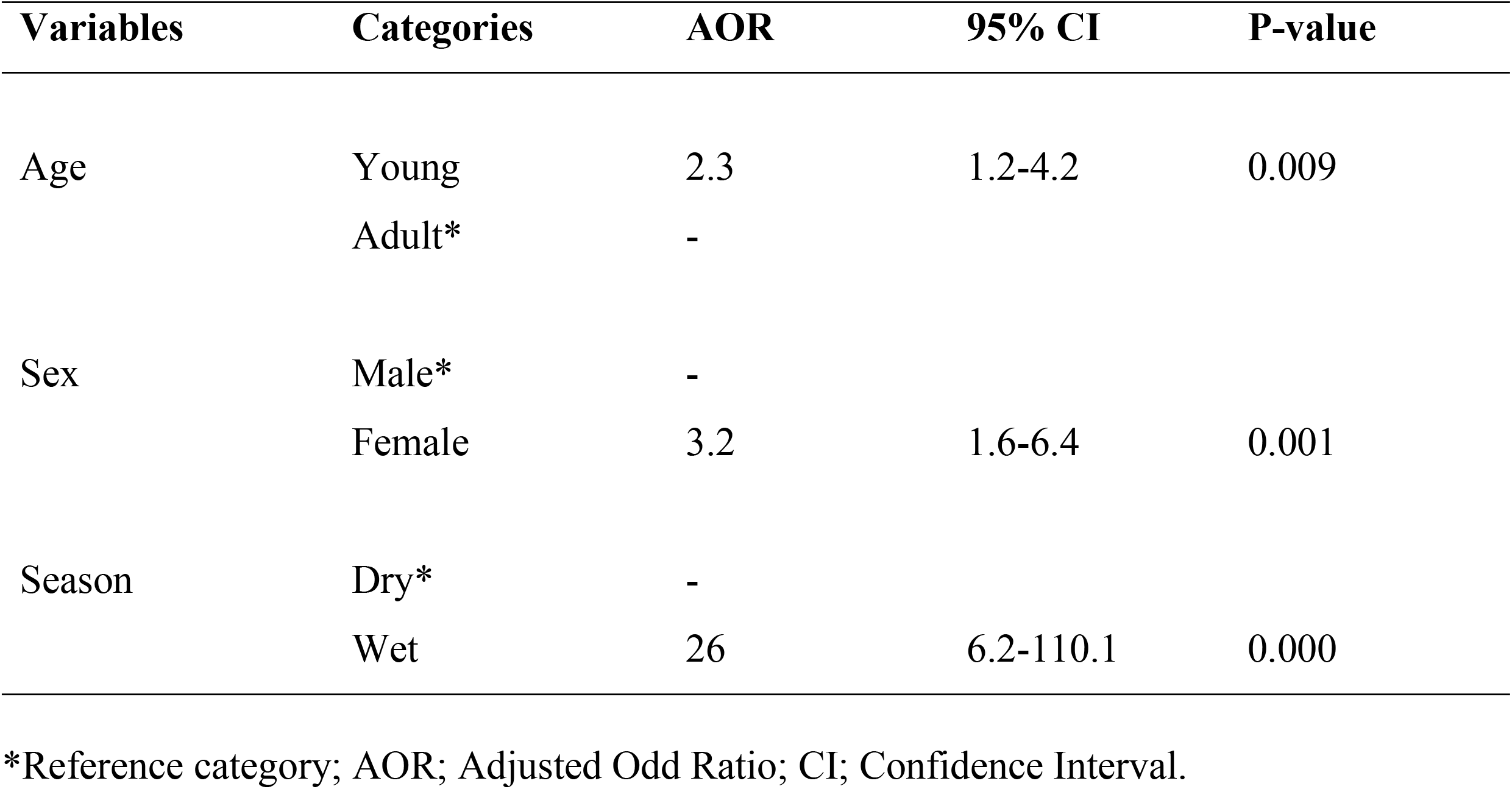
Final model on multivariable analysis of risk factors related to camelpox occurrence.

## Discussion

Out of the 374 dromedary camels sampled from two districts within the Jarar Zone, 60 tested positive for camelpox using a sandwich ELISA test. This indicates a high prevalence of the disease in the examined camel herds. The purpose of this study was to conduct a sero-epidemiological investigation of camelpox in order to estimate the prevalence and identify the associated risk factors in the study area.

This result agrees with a previous report of camelpox from a study conducted in the Afar pastoral region. The previous study reported a camelpox seropositivity of 19.3% using the virus neutralization test, which detected the presence of viral antibodies in dromedary camels investigated by Aregawi *et al*. [24]. However, our results differ from those obtained in previous studies. These previous studies showed a higher prevalence in the Amibara district of the Afar region, with a respective prevalence of 21.6% [24]. Additionally, our findings contradicted another study conducted by Ayelet et al.[25], which reported a lower prevalence of 4.50% in the Afar Region. These differences could be due to variations in agro-geographical location, sample size, and animal herding practices.

In the current study area of the Somali region, there is a strong clan-based segregation of animals and use of rangelands that restricts disease transmission among camel herds. In contrast, in the Afar Region, mixing of animals from different areas is common at communal grazing and watering areas [26]. The observed higher prevalence of camelpox in this study compared to other studies from different parts of Ethiopia, based on clinical cases reporting prevalence rates of 3%, 14.2%, and 3.8% respectively [27, 28], is not unexpected. Serological prevalence is often higher than clinical prevalence, as poxvirus antibodies can be detected in animal sera much more frequently than poxviruses can be isolated from clinical cases [29]. Similar findings were also reported for other camel diseases, such as trypanosomosis, where parasite antibody prevalence was observed to be far higher than the prevalence of live parasites [30].

Age, sex, and season were identified as statistically significant factors associated with occurrences of camelpox and camelpox antibody seropositivity. The study found that seroprevalence was significantly higher in young dromedaries than in adults (p<0.05). This finding can be explained by the fact that younger camels have less developed immune systems, making them more susceptible to infection and resulting in higher seropositivity rates. Another possible explanation is the decrease in maternal antibodies and increased exposure to the virus, which leads to higher susceptibility to camelpox infection in young camels compared to adults [31]. On the other hand, older camels may have acquired immunity through previous exposures or vaccinations, resulting in lower seropositivity [32]. This study contradicts a previous study conducted in the Afar region, which showed a significantly higher seroprevalence of camelpox antigen in adult age groups (>4 years) compared to younger age groups (>6 months and <4 years) [33, 34]. This discrepancy could be due to the unequal proportions of the two age groups during sampling. Additionally, the variation in the severity of clinical signs may reflect differences between the strains of CMLV or the animals sampled, such as old camels (>10 years old), which are known to have poor immunity [35].

Based on sex, the current study found that females had a higher prevalence of camelpox antibodies, accounting for 21%. This difference was statistically significant (p<0.05). The findings suggest that hormonal influences, such as higher levels of prolactin and progesterone, may affect immune responses in females. This could potentially make them more susceptible to camelpox infections and hinder their ability to fight off or control the disease [36]. Other factors, such as pregnancy and lactation stress, may also increase the susceptibility of female camels to camelpox. Additionally, the breeding behavior of pox-infected males could contribute to the spread of the disease to multiple females. However, the role of sex in susceptibility to camelpox remains disputed in the scientific literature. Other studies have produced conflicting results on this aspect [37, 38].

This study revealed a seasonal variation in the occurrence of camelpox. The disease was found to be more common during the wet rainy season (27.0%) compared to the dry season (1.0%), and this difference was statistically significant (p-value<0.05). This finding aligns with a previous study conducted in Ethiopia’s Afar pastoral region by Weldegebrial et al. [39]. They also reported that camelpox occurs frequently during the minor and major rainy seasons and described the disease as endemic in the region. Similarly, other studies have shown that camelpox outbreaks increase during the rainy season. In these cases, a more severe form of the disease occurs during wet seasons, while a milder form is seen during dry seasons. One possible explanation for this pattern is that wet seasons may provide more favorable conditions for virus replication or enhance the survival of virus particles outside the host. Consequently, seropositivity rates for camelpox may be higher during these periods. Another possible reason for this seasonal difference is that camelpox is primarily transmitted by arthropod vectors, such as mosquitoes or ticks, and the abundance of these vectors often varies with seasonal changes. Consequently, increased vector activity during wet seasons can result in a higher transmission rate of the camelpox virus, leading to increased camelpox seropositivity [34].

## Conclusion

In conclusion, our study aimed to investigate the epidemiology of camelpox in selected districts of the Jarar Zone in the Somali Region of Ethiopia. Through a sero-epidemiological investigation, we estimated the prevalence of camelpox and identified the associated risk factors contributing to the occurrence of the disease. Our findings revealed a significant presence of camelpox antibodies circulating within the examined camel herds in both districts. This highlights the importance of implementing effective disease management and control measures to minimize the impact of camelpox on camel populations and reduce economic losses. By identifying significant hypothesized risk factors such as age, sex, and seasons, we gained valuable insights into the transmission dynamics of camelpox. Therefore, our study contributes to the understanding of camelpox epidemiology in the study area and provides a solid foundation for proactive disease management strategies. These strategies should include timely implementation of vaccination campaigns, improved husbandry practices, awareness programs, and the provision of useful information to stakeholders to prevent and control the spread and burden of the disease. Continued surveillance and further research are warranted to explore the genetic diversity of camelpox viruses circulating in the region.

### List of abbreviations

AOR: Adjusted Odds Ratio
CI: Confidence Interval
PDS: Participatory Disease Surveillance
CMLV: Camelpox Virus
VNT: Viral Neutralization Test
JRLDC: Jigjiga Regional Laboratory Diagnostic Center
OD: Optical Density
sELISA: Sandwich Enzyme-Linked Immunosorbent Assay.

## Author contributions

Hassan Abdi Arog and Abdullahi Adan Ahad: designed the study and collected the data; Abdullahi Adan Ahad: performed the data entry; Haben Fesseha and Hassan Abdi Arog: analyzed and interpreted the data; Abdullahi Adan Ahad: drafted the original manuscript; All authors: contributed to the writing and approval of the paper.

## Funding statement

This research did not receive any external funding.

## Availability of data and materials

The data set generated and analyzed during the current study are not publicly available but can be obtained from the corresponding author upon a reasonable request.

## Acknowledgements

The authors would like to acknowledge the Jigjiga Regional Veterinary Laboratory and Diagnostic Center (JRVLDC) for aiding with materials used in the study, sample preparation, and analysis. The authors also extend their gratitude to the camel owners in the study area for their willingness and cooperation.

## Conflicts of interest

The authors have stated that they do not have any competing conflicts of interest.

---

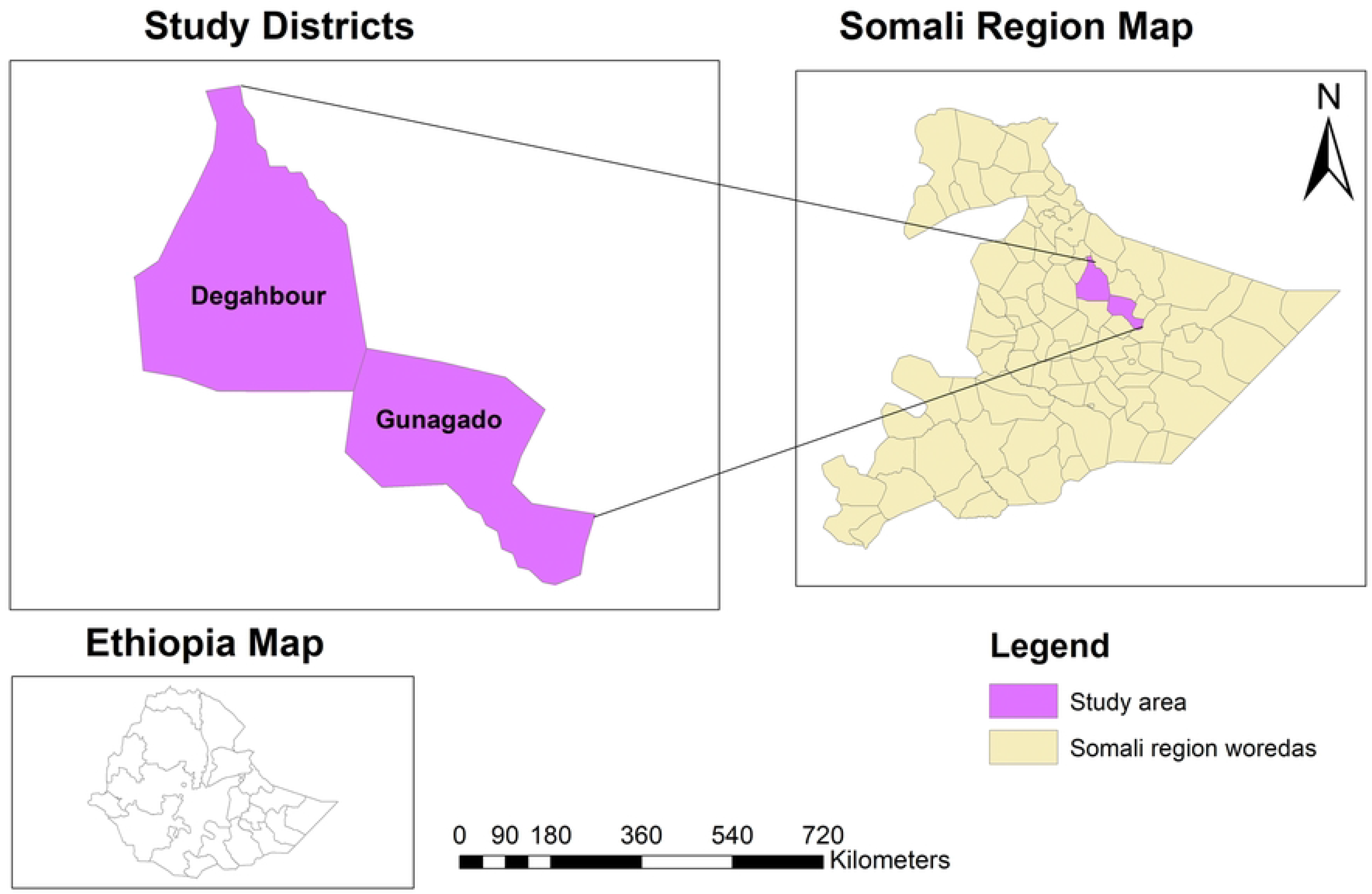

## Notes

### Competing Interest Statement

The authors have declared no competing interest.

